# Mechanisms of Synchrony in Heterogeneous Inhibitory Interneuronal Networks for Type 1 versus Type 2 Excitability

**DOI:** 10.1101/802330

**Authors:** Ruben A. Tikidji-Hamburyan, Carmen C. Canavier

## Abstract

PV+ fast spiking basket interneurons are often implicated in gamma rhythms. Here we focus on mechanisms present in purely inhibitory networks. Neurons with type 1 excitability can fire arbitrarily slowly, whereas those with type 2 excitability cannot fire below a minimum frequency. We systematically examine how excitability type affects synchronization of individual spikes to a population rhythm in the presence of heterogeneity and noise, using model neurons of each type with matched F/I curve, input resistance, time constant and action potential shape. Population synchrony in noisy heterogeneous networks is maintained because neurons either fire within a tight time window or skip that cycle. Type 2 neurons with hyperpolarizing inhibition skip cycles due to their intrinsic dynamics; we show here the cycle skipping mechanism for type 1 neurons or type 2 neurons with shunting inhibition is synaptic and not intrinsic. Type 2 neurons are more resistant than type 1 to partial and complete suppression in networks with hyperpolarizing inhibition that exhibit network gamma. Moreover, type 2 neurons are recruited more rapidly and more completely into theta-nested gamma. In contrast, type 1 networks perform better with shunting inhibition on both counts, because the nonlinear dynamics in that case favor suppression of type 2 compared to type 1 neurons. Conductances that control excitability type may provide a therapeutic target to improve spatial and working memory and other tasks that rely on gamma synchrony or phase amplitude coupling.

**Author Summary:** The collective, synchronized activity of neurons produces brain rhythms. These rhythms are thought to subserve cognitive functions such as attention and memory encoding and retrieval. Faster rhythms are nested in slower rhythms as a putative way of chunking information. A subset of neurons called fast spiking basket cells tend to inhibit other neurons from firing. These neurons play an important role in oscillations, and in the coupling of faster oscillations to slower ones. In some brain regions these neurons can fire arbitrarily slowly (type 1 dynamics) whereas in others they cannot fire below a minimum cutoff frequency (type 2 dynamics). Mathematically, these distinct origins of rhythmic firing are signatures of very different dynamics. Here, we show that these distinct excitability types affect the ability of networks of these neurons to synchronize their fast oscillatory activity, as well as the ability of slower oscillations to modulate these fast oscillations. The exact nature of the inhibitory coupling, which may vary between brain regions, determines which type synchronizes better and is modulated better.

## Introduction

PV+ fast spiking (FS) basket interneurons are often implicated in gamma rhythms [1–5], which in the hippocampus are thought to organize information for memory encoding and retrieval [6–8]. Gamma frequency (30–80 Hz) oscillations are thought to serve as substrates for working memory, conceptual categorization, and attention [9], and are altered in psychiatric disorders, for example schizophrenia, dementia, and autism [10]. PV+ FS basket cells play an important role in theta nested gamma [11]. Moreover, nesting of gamma within theta has been proposed a substrate for episodic memory [12] and disruption of theta nested gamma has been proposed to explain deficits in spatial memory in temporal lobe epilepsy [13, 14].

Here we focus on mechanisms present in purely inhibitory networks, and the role of excitability type of the FS interneurons in gamma frequency synchronization and cross frequency synchronization with theta oscillations. Neurons with Hodgkin’s type 1 excitability are able to spike arbitrarily slowly, whereas those with type 2 excitability have an abrupt onset of repetitive firing that cannot be maintained below a threshold frequency [15, 16]. Our recent work [17] shows that FS neurons likely exhibit type 2 excitability in the medial entorhinal cortex, and others have shown that they exhibit type 2 excitability in striatum [18] and neocortex [19]. However, previous work supports type 1 excitability in hippocampal area CA1 [20–22]. We will focus on reduced, 2D models here. The advantage of 2D models is that they can be visualized in a two-dimensional phase plane that reveals the essential features of the intrinsic dynamics. For type 2 excitability, there is a circular flow near the associated subcritical Hopf bifurcation that results in subthreshold oscillations, postinhibitory rebound inherent in the spike generating mechanism, and a resonant spiking frequency. These features are absent in neurons with type 1 excitability. In contrast, type 1 neurons rely on a saddle-node-on-an-invariant-circle (SNIC) bifurcation, which creates a slow channel in the phase plane where the net current is very small, allowing the neurons to spike arbitrarily slowly.

Previously [17], we noted that cycle skipping was a way for a network to preserve synchrony of individual spikes with the population in the presence of heterogeneity and noise. In other words, spikes that would have occurred too late to be in synchrony with the population are suppressed. In the case of type 2 excitability, this can be accomplished via the intrinsic dynamics, because in all regimes (excitable, bistable and oscillatory) there is a boundary between trajectories that lead to spiking and those that lead to cycle skipping. However, we did not calibrate the models we compared in such a way as to maximize the fairness of the comparison. Moreover, we only looked at hyperpolarizing and not shunting inhibition.

Fast, ionotropic inhibition in the central nervous system in generally mediated by GABA_A_ receptors, with chloride ions as the charge carrier. The reversal potential of these channels depends on the intracellular concentration of chloride. For quiescent neurons, for hyperpolarizing inhibition the synaptic reversal potential that is negative to the resting potential, whereas for shunting inhibition the reversal potential is more positive than the resting potential but negative to the spike threshold. For oscillatory neurons, the membrane potential during the interspike interval substitutes for the resting potential [23]. Shunting inhibition is sometimes defined as an increase in synaptic conductance in the absence of an obvious change in membrane potential [24]. In our model neurons, a reversal potential of −75 mV produces hyperpolarization whereas −65 mV does not, so we classify the latter as shunting in our context. The reversal potential for chloride in fast spiking basket cells *in vitro* is about −52 mV [25] in the dentate gyrus, which implies shunting inhibition in those cells. Previous studies in CA1 and CA3 concluded that inhibition between basket cells was hyperpolarizing [26, 27]. However, the Cl-concentration can be modulated [28], and there can be intracellular gradients in chloride concentration [29]. Therefore, there is sufficient uncertainty regarding the precise reversal potential of chloride at synapses between interneurons to warrant a systematic study of both types of inhibition.

Here, we systematically examine how excitability type affects synchronization of individual spikes to a population rhythm in the presence of heterogeneity and noise. Simple 2D model neurons of each type were calibrated to have a very similar F/I curve, input resistance, time constant and action potential shape in order to isolate the consequences of excitability type alone on the robustness of synchronization in inhibitory interneuronal networks.

## Results

### Phenomenological model of type 1 and type 2 excitability

We began with the standard 2D reduction [30] of the 4D Hodgkin-Huxley model [31]. The activation of the voltage-dependent sodium current was assumed to be fast and set to its steady-state value with respect to the membrane potential (*m*_∞_ (*v*)). The inactivation variable for the voltage-dependent Na^+^ current (*h*) was yoked to the variable (*n*) for the activation of the delayed rectifier K^+^ current, under the assumption that the slow time scale for these variables was similar. This 2D system with one fast variable (membrane potential, *v*) and one slow variable (*n*) is amenable to phase plane analysis under fast/slow assumptions [32]:

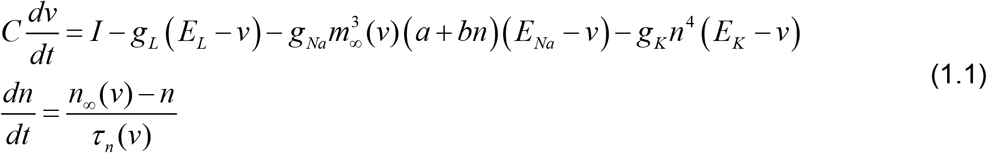

where: 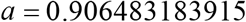 and b 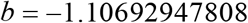 are the offset and slope for the 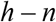 linear regression; 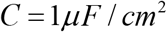 is a membrane capacitance; *I* is applied current in 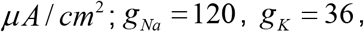 are *g_L_* conductances in *mS / cm*^2^ for sodium, potassium, and leak currents with corresponding reversal potentials 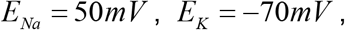 and 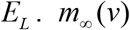 and *n*_∞_(*v*) are steady state activations for sodium and potassium channels; and 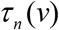 is the time constant for potassium channel activation in ms:

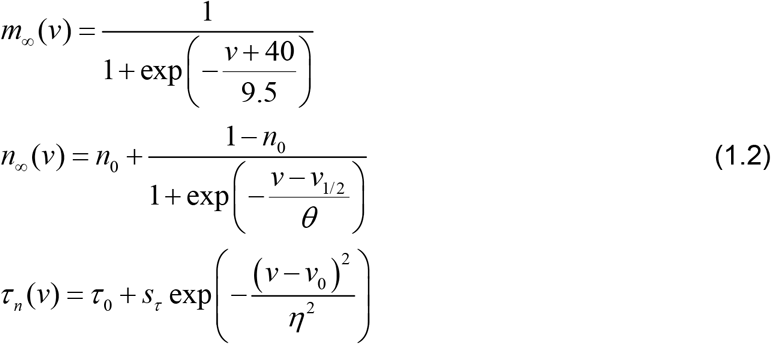

Figure 1 shows a phase plane analysis of the bifurcations underlying type 1 (Fig. 1A) and type 2 (Fig 1B) excitability. In the phase plane, fixed points occur at the intersection of the n-nullcline (gray curve), which is simply the steady state activation curve for the *n* activation variable, and the voltage nullcline at rest (red curve), which is the set of values of *n* and *v* for which the net ionic current plus any applied current is zero. The voltage nullcline is N-shaped with three branches, but in order to emphasize the bifurcation point, only the stable left and unstable middle branch and their associated fixed points are shown in Fig 1. For this fast/slow system the leftmost branch is stable, and the middle branch is unstable due to the autocatalytic sodium current. At rest (with no applied current), the fixed point (filled circle) on the left branch is stable and determines the resting potential at −68 mV for both the type 1 (Fig. 1A1) and type 2 (Fig. 1B1) cases. The first important difference is that for type 2 there is a single fixed point at all values of applied current, whereas for type 1 the number of fixed points depends upon the level of the applied current.

**Figure 1.**
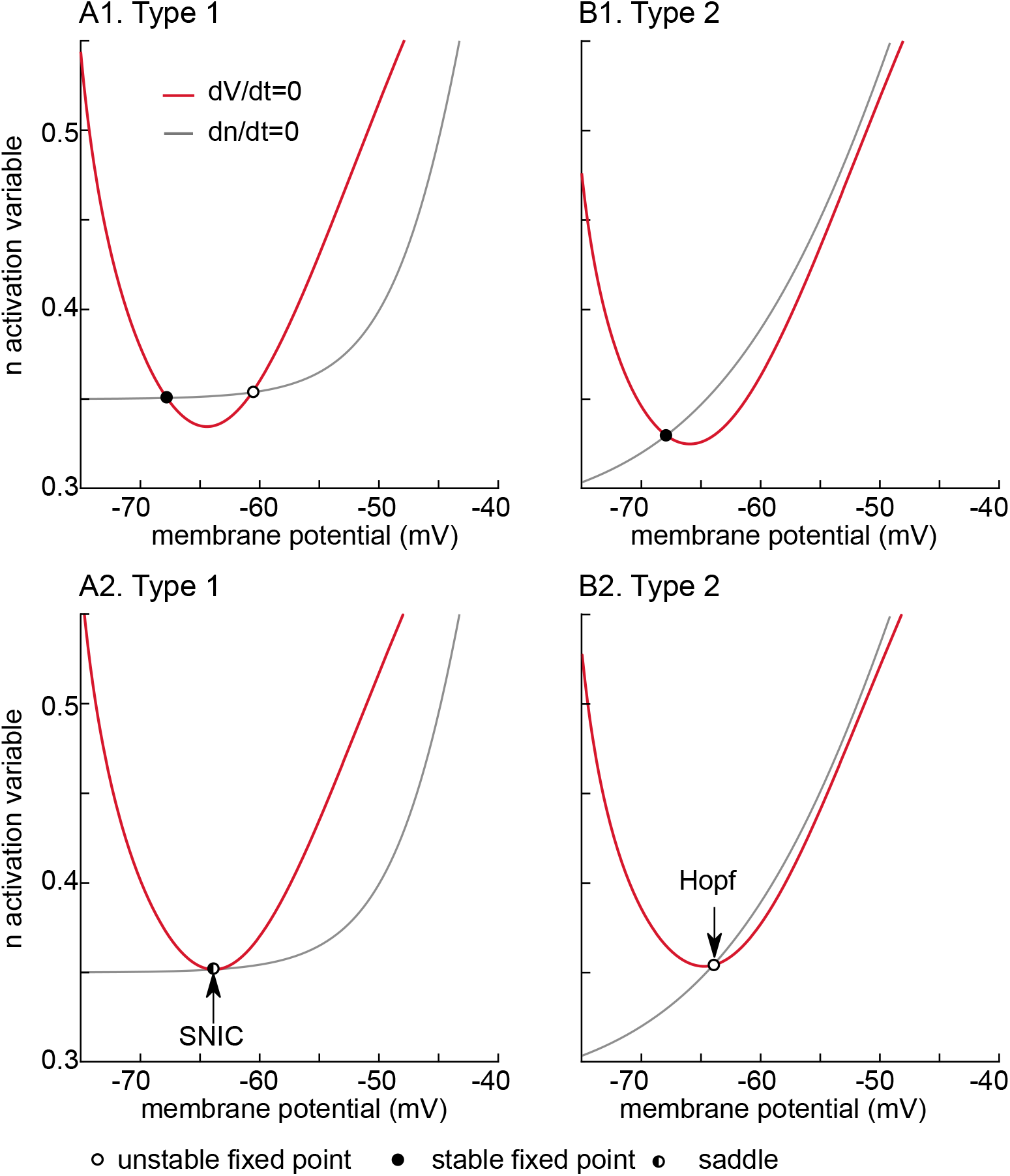
Phase plane analysis of model bifurcation structure. Red curves indicate the left-most two branches of the cubic V-nullcline on which the rate of change of membrane potential is zero. The gray curve is the n-nullcline on which the rate of change of the slow variable is zero. Filled and open cycles are stable and unstable fixed points. The half filled circle is the saddle node. Under the assumption of fast/slow dynamics, fixed points on the left branch are stable and those on the other branch shown are unstable. A. Type 1. A1. Quiescent neuron with no bias current has a stable resting potential (filled circle) on the left branch at −68 mV. Only one of the two fixed points on the unstable branch appear in this view, chosen to emphasize the bifurcation. A2. The n-nullcline is tangent to the V-nullcline as a stable and unstable fixed point collide to form a saddle node with *I* = 1.38*μ A / cm*^2^. The bifurcation is only a SNIC if a limit cycle is born simultaneously and emanates from the saddle node. B. Type 2. B1. Quiescent neuron with no bias current has a stable resting potential (filled circle) on the left branch at −68 mV. There is only a single fixed point. B2. The Hopf bifurcation occurs as the fixed point loses stability as it moves onto the unstable branch at *I* = 2.11*μ A / cm*^2^.

For type 1 at rest (Fig. 1A1), in addition to the stable fixed point that determines the resting potential, there is an unstable fixed (open circle) point on the middle branch, as well as another one on the unstable branch that is not shown. As the applied current is increased to 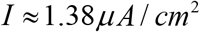 the stable and unstable fixed points shown collide (Fig. 1A2) and form a saddle node (circle half filled). This collision coincides with the birth of limit cycle surrounding the rightmost unstable fixed point on the unstable branch (not shown), therefore the onset of spiking occurs at a saddle-node-on-an-invariant-limit-cycle (SNIC) bifurcation [16, 33]. As the applied current is increased further and the colliding fixed points annihilate each other. The neuron is left with no stable resting potential. The intersection of the limit cycle with the saddle node gives rise to a trajectory with infinite period. As the applied current is increased further, the trajectory must pass through a gap between the two nullclines; the rate of passage though this bottleneck is arbitrarily slow with increasing proximity to the bifurcation, hence the arbitrarily slow frequencies obtainable by type 1 neurons shown in Fig. 2A (black dots).

**Figure 2.**
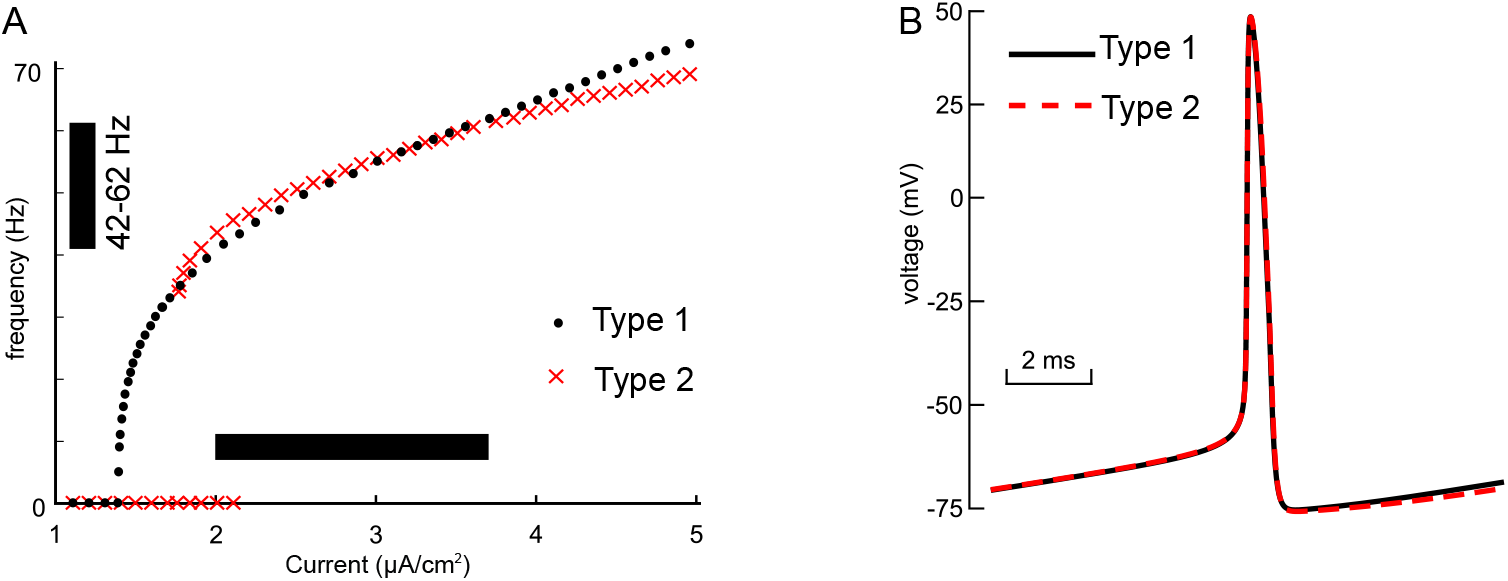
Model Calibration. The parameters of the model for both types (parameters given in Table 1, attributes in Table 2) were adjusted to make the comparison as fair as possible. A. Frequency/ current (f/I) curves for type 1 (black dots) and type 2 (red crosses) overlap for the range of bias currents (horizontal bar) used in our heterogeneous networks. The vertical bar shows the range of frequencies exhibited at the heterogeneous bias current levels. The f/I curves were measured using current steps of sufficient duration to allow any transients to die out and establish a steady frequency. Arbitrarily slow frequencies can be obtained for type 1 (not shown) near the bifurcation, but type 2 has a minimum cutoff frequency below which it cannot fire. Between current steps, the membrane potential was returned to its resting value. The bistable range for type 2 is evident from the current values at which a zero frequency quiescent solution coexists with a repetitively firing solution. This region was determined using XPPAUTO. B. The spike shapes and the interspike interval are very similar for type 1 (black curve) and type 2 (red dashed curve). I = 2.85*μ* A / cm^2^

**Table 1.** Parameters for Type-I and Type-II excitabilities of the neuron model

| Type | $g_L(mS/cm^2)$ | $E_L(mV)$ | $n_0$ | $v_{1/2}(mV)$ | $\theta(mV)$ | $\tau_0(ms)$ | $s_\tau(ms)$ | $v_0(mV)$ | $\eta(mV)$ |
| --- | --- | --- | --- | --- | --- | --- | --- | --- | --- |
| I | 0.3 | -54.3 | 0.35 | -40.0 | 4.0 | 0.46 | 3.5 | -60.5 | 35.9 |
| II | 0.1 | -39.0 | 0.28 | -44.5 | 9.0 | 0.5 | 5.0 | -60.0 | 30.0 |

**Table 2.**
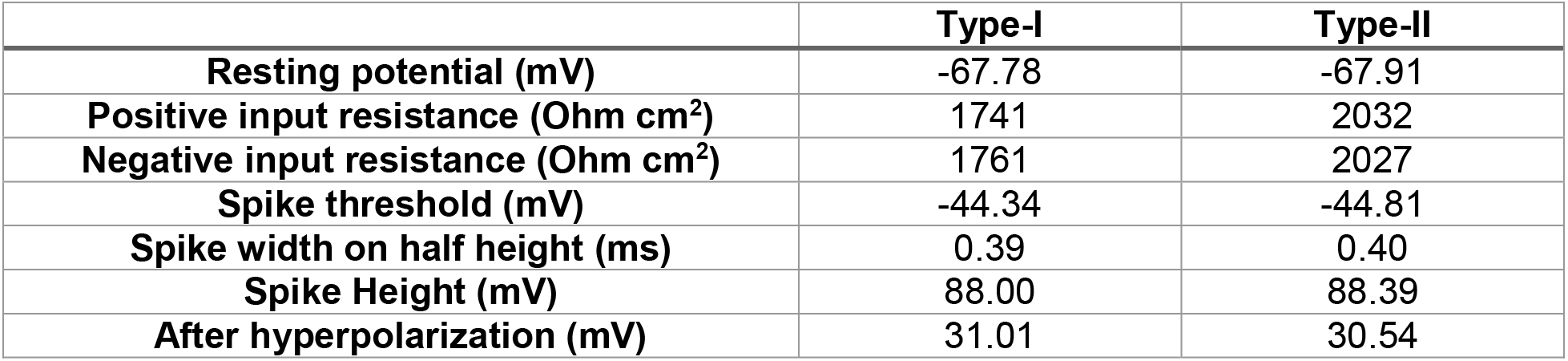
Characteristics of membrane passive properties and spike shape in Type-I and Type-II modes

In contrast, for type 2 excitability the Hopf bifurcation occurs when the lone stable fixed point corresponding to the rest potential loses stability (Fig. 1B1). When the applied depolarizing current reaches 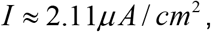 the fixed point (Fig. 1B2, open circle) moves past the trough of the nullcline onto the unstable branch of the voltage nullcline. The Hopf bifurcation is subcritical [16,33,34] because in the range of input currents between 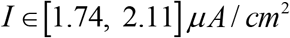 the model exhibits bistability between stable limit cycle corresponding to repetitive spiking and a stable fixed point (Fig. 2A, overlap in the red cross at 0 frequency and at a nonzero frequency). These attractors are separated by unstable limit cycle (not shown). The stable and unstable limit cycles collide and annihilate each other in a saddle node of periodics at 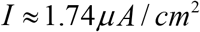. Excitability type is defined as the response of a quiescent neuron as the applied current is increased [15]; the sudden onset of spiking as the applied current is increased occurs at about 30 Hz when the quiescent state loses stability as shown in Fig. 1B.

Model parameters 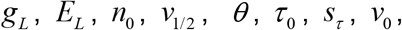 and *η* are different for Type-I and Type-II regimes and are given in Table 1. These parameters were adjusted for the two excitability type in order to keep the input resistance 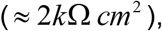 time constant 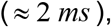 F/I curve (Fig. 2A) and spike shape (Fig. 2B) as similar as possible. The shape of the steady state activation curve for the delayed rectifier is non-physiological because it does not go to zero. This phenomenological compromise was necessary to keep the input resistance of quiescent model neurons comparable and does not affect the dynamics of interest near the bifurcations.

### Steady State Synchrony in Oscillatory Networks

We constructed networks of 300 sparsely and randomly connected neurons with heterogeneity in frequency by distributing the bias current uniformly along a region of the F/I curve that spanned a 20 Hz range (see black bars in Fig. 2A). The conduction delays between neurons were also uniformly distributed between 0.7 and 3.5 ms. In our previous work [35] in a homogeneous network, delays on the short end of this range favor a solution with two subclusters in antiphase, whereas delays at the longer end of the range favor global synchrony of a single cluster. Using a mixture of delays results in solutions that are not obviously one or two clusters, but are transitional between these two extremes.

### Steady State Synchrony in Oscillatory Networks-Hyperpolarizing Inhibition

Figure 3A and B top left show representative traces from network simulations with hyperpolarizing inhibition showing how individual neurons of both types in noisy, sparsely connected and heterogeneous networks skip random cycles. However, neurons in both populations remain synchronized with the population oscillation when they do fire, which is evident from raster plots Fig. 3A and B bottom-left. The cycle skipping is evident in the histograms of interspike intervals (ISI) (Fig. 3A and B bottom right) across all neurons. There are peaks at the network period and at integer multiples of the cycle period, corresponding to how many cycles were skipped during that interval. Neurons in the raster plots are sorted by bias current with the fastest firing cells at the top. This example shows that the slower firing neurons at the bottom of the raster plots have a much greater likelihood of being completely or partly suppressed for type 1 networks compared to type 2 networks. The histogram of the average number of network cycles each individual neuron participated (top right, Fig. 3A and B) in is much flatter for type 1 than type 2, because the type 2 histogram is skewed to higher participation rates.

**Figure 3.**
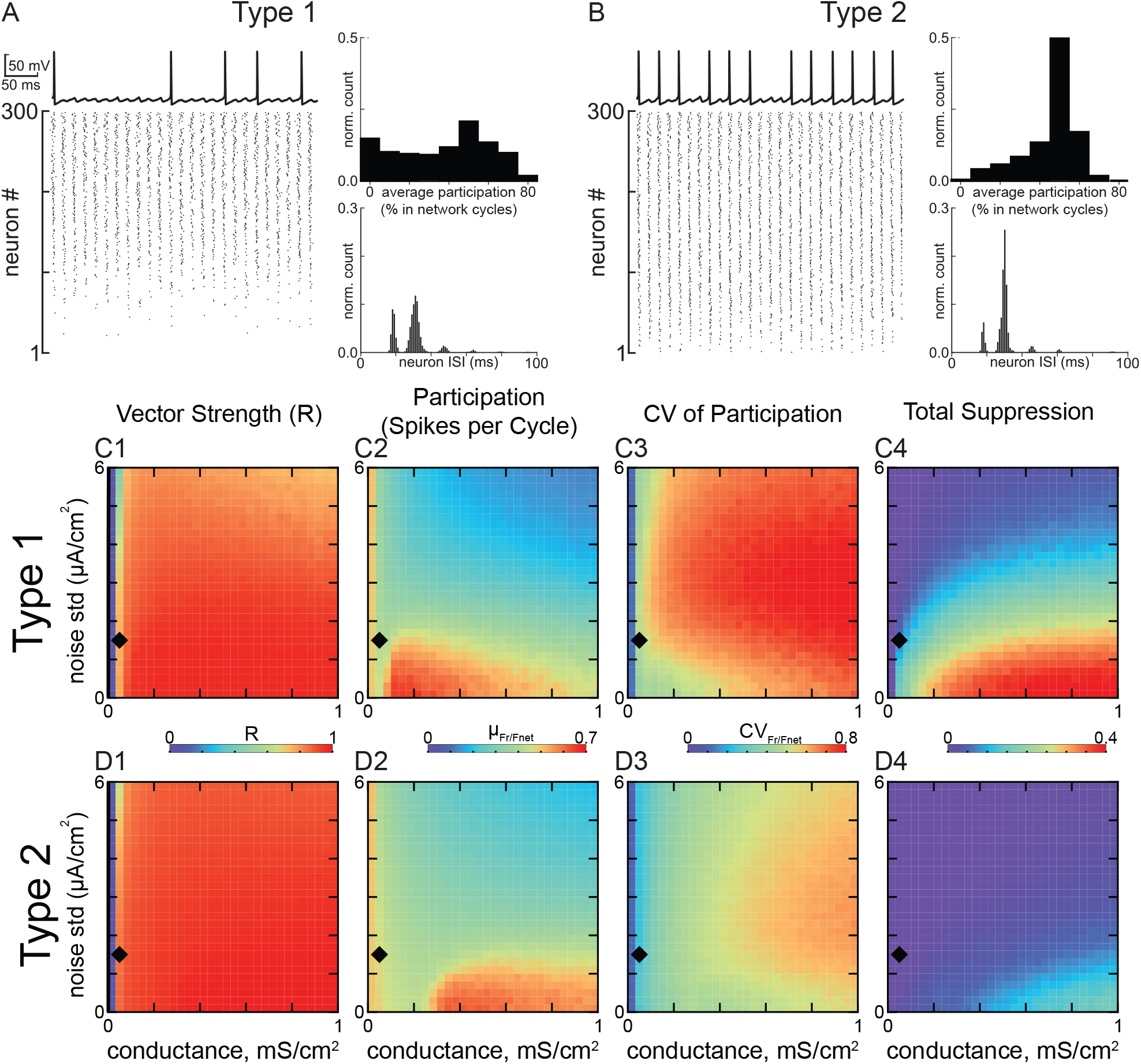
Steady State Synchronization of Heterogeneous Inhibitory Interneuronal Networks with Hyperpolarizing Inhibition. A. Networks of type 1 neurons with synaptic conductance 0.05 mS/cm^2^ and noise standard deviation 1.5 µA/cm^2^, indicated by black diamonds in C and D. Bottom left: raster plot for 300 neurons, ordered by the level of applied depolarizing bias current, with the most depolarized neurons at the top. Top left: Representative membrane potential trace for an individual neuron that clearly exhibits a subthreshold oscillation due to network activity and cycle skipping. Bottom right: Histogram of interspike intervals across the population. Top right: Histogram of average participation for all neurons in the network. B. Networks of type 2 neurons with the same parameters as in A. The four panels are the same as in A. C. 2D parameter sweep in conductance strength and noise standard deviation for type 1 networks. Heatmaps from left to right: C1: vector strength R of synchronization of individual spikes with the network oscillation, C2: average participation for neurons that are not totally suppressed calculated as the mean µ_Fr/Fnet_ of the frequency of spiking neurons normalized by the population frequency, C3: the coefficient of variation of the participation CV_Fr/Fnet_, and C4: the fraction of completely suppressed neurons. D. 2D parameter sweep for type 2 networks. Heatmaps are the same as in C.

This tendency for greater suppression of type 1 neurons with hyperpolarizing inhibition was preserved across a large range of the 2D parameter space of synaptic conductance strength and standard deviation of the additive Gaussian current noise, as shown by the heatmaps in Figs. 3 C and D. The diamond in the heatmap indicates the parameters used to generate the raster in panels A and B. The leftmost heatmap gives the vector strength measure of population synchrony. For both Type 1 (Fig 3C1) and Type 2 (Fig 3D1), a minimum amount of conductance is required for synchrony (blue strip at the left is unsynchronized). Both networks perform fairly well for synchrony of spikes with the population rhythm over this range, although Type 2 performs a little better at high noise levels and stronger conductance. The vector strength is quite high, and exceeds 0.8 almost everywhere. The average spikes per cycle (excluding neurons that are completely suppressed) decreases with increasing noise, but again similar for the two types of networks (Fig 3C2 and D2). The major difference is evident in the heatmaps (Fig. 3C3 and D3) for the coefficient of variation (CV) for participation across the population. As expected from the participation histograms shown in panels A and B (top right of each), the CV is greater for Type 1, reflecting the greater abundance of partially suppressed neurons. The CV is in the range 0.5-0.8 for type 1compared to 0.3-0.6 for Type 2. This discrepancy would be even greater had we included the factions of totally suppressed neurons illustrated in the rightmost histograms(Fig. 3C4 and D4), which clearly show a far greater fraction of totally suppressed neurons for type 1. Stronger noise desynchronizes and stronger inhibition promotes suppression, so outside the regime of interest neither type performs well and the difference between type 1 and 2 fades.

**Figure 4.**
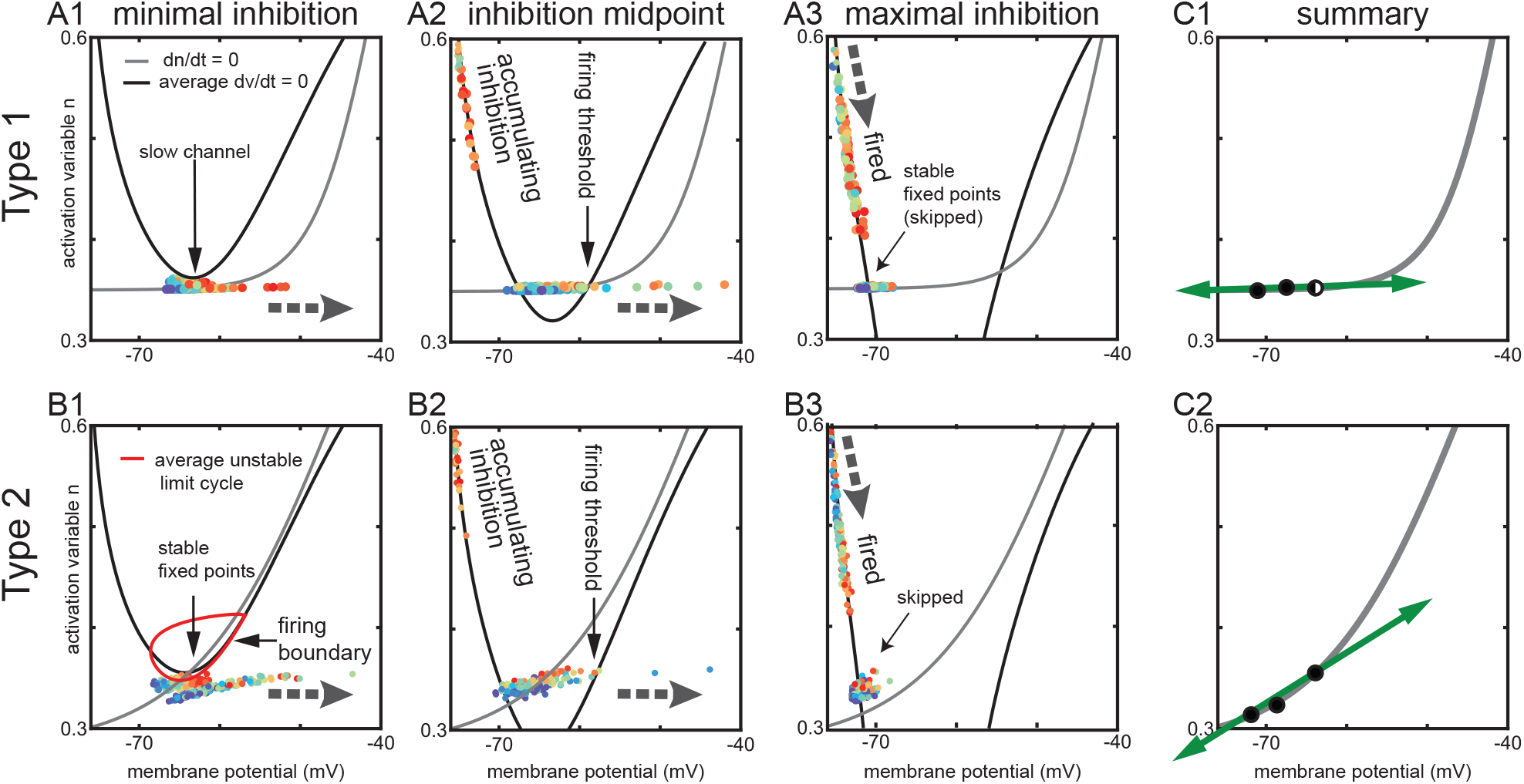
Phase Plane Analysis of Heterogeneous Inhibitory Interneuronal Networks with Hyperpolarizing Inhibition. Large gray dashed arrows show the local direction of motion in the vector field. Other thin black arrows simply point out a specific feature. These phase plane portraits are similar to those in Fig. 1 with two exceptions. First, the V-nullcline (black curves) shown is an average across the heterogeneous population. Thus, the fixed points shown are also an average, and actually differ for each neuron. Second, the time-varying levels of inhibition between the neurons causes the position of the average V-nullcline to fluctuate in time. The slow n-nullcine (gray) is constant for each type. Each dot represents the current position of one neuron in the network, with the fastest neurons in red and the slowest in blue. Movement in the horizontal direction is fast compared to that in the vertical direction because the slow variable is plotted along the y axis. A. Type 1. A1. Minimal inhibition creates a slow channel between the nullclines (diagonal arrow). A2. Inhibition midpoint. As neurons escape to the right (horizontal arrow shows direction of motion) and fire an action potential, inhibition accumulates and moves the average V-nullcline down. The right branch of the V-nullcline is the firing threshold (vertical arrow). A3. Maximal inhibition. The population is segregated into two groups. One group that fired on the most recent network cycle slowly moves down (thick arrow at top shows direction of motion) along the stable left branch of the average V-nullcline. The other group (diagonal arrow) skipped this cycle and tends to line up along the flat portion of the n-nullcline. B. Type 2. B1. Minimal inhibition. In contrast to the absence of a fixed point in A1, there is a stable fixed point (technically fixed points, vertical arrow) at the intersection of the average V-nullcline and the n-nullcine. This fixed point is surrounded by an unstable limit cycle (red curve) that forms the boundary between spiking and quiescent trajectories. B2. Inhibition midpoint. As in A2 inhibition accumulates as neurons escape to the right and fire action potentials. The right branch of the Vnullcline is now the firing threshold (vertical arrow). B3. Inhibition maximum. As in A3, two groups are evident corresponding to those that fired and those that skipped. C. Summary. C1. This figure shows that the movement of the fixed point as inhibition waxes and wanes is (leftward) in the fast, horizontal direction for type 1. C2. In contrast, the downward and leftward movement of the fixed point for type 2 has a component in the slow direction, which helps equalize the opportunity of slow neurons to fire as compared to fast neurons.

The 2D phase plane representation of network activity shown in Figure 4 illustrates the mechanism underlying greater suppression of type 1 compared to type 2 neurons in networks with hyperpolarizing inhibition. The n-nullcline for the slow variable (gray curve) was constant in time and across the population. On the other hand, the membrane potential V-nullcline in the absence of inhibition was different for every neuron because of the heterogeneity in applied bias current. This required averaging the V-nullcline across the population. Moreover, the level of inhibition is a third state variable per neuron, and the average level of inhibition in the network varies in time. Therefore, the 2D phase plane representation of the V-nullcline (black curve) actually constitutes a movie (supplemental material) and snapshots of this portrait are given for minimal, half-amplitude, and maximal inhibition in Fig. 4 A1-3 and B1-3 for type 1 and 2, respectively. Each dot represents the current position of a neuron in this phase space, with the neurons with the highest depolarizing bias current shown in red and those with the least in blue.

At minimum (but nonzero) inhibition, most neurons are in a near threshold regime (Fig. 4 A1 and B1). The average nullcline portrait is close to the SNIC bifurcation. For type 1 networks: a slow channel (arrow) between the two nullclines arises as the system approaches the SNIC bifurcation (see Fig. 1A2). By definition, the rate of change of a variable is zero on its nullcline, and very near the nullcline, the rate of change is quite slow. In the channel, the rate of change of both variables is quite slow, but the direction of the rate of change of membrane potential is positive, enabling arbitrarily slow firing rates. Moreover, the level of bias current imposes order on the trajectories, with the fastest cells positioned nearest the firing threshold. As neurons escape from the channel and fire action potentials, inhibition accumulates as the inhibitory post-synaptic potentials (IPSPs) from the spiking neurons (shown moving down the left branch of the V-nullcline) summate in Fig. 4A2. The accumulating inhibition moves the average V-nullcline down and to the left, closing the channel and creating two fixed points (as in Fig. 1A1) at the intersection with the n-nullcline. The unstable branch of the V-nullcline (arrow indicating firing threshold points to this branch) separates neurons into two groups. Neurons whose trajectory has already moved to the right of this branch will fire an action potential, but those that fall on the left side of this boundary when the channel closes will skip this network cycle. In general, the neurons that skip are the slower neurons; they will move leftward towards the stable fixed point. The two groups are clearly shown in Fig. 4A3 at the point of maximal inhibition after all spiking neurons have fired on a given cycle. The group labeled fired is recovering from the after-hyperpolarizing potential (AHP) along the stable V-nullcline branch. The group labeled skipped is trapped on their fixed point and did not participate in the previous cycle. The intersection of the average nullclines is the average fixed point; the actual fixed point for the slower neurons lies to the left of the average and for the faster neurons it lies to the right. Since the connectivity is random, some neurons receive more inhibitory inputs than others, which adds a leftward bias to their fixed point. Both groups are aligned according to the net current due to a combination of the bias current and the number of active inhibitory inputs. Thus, the composition of the suppressed population on each cycle is influenced by the number of active inhibitory inputs. When the maximal inhibition decays, the phase portrait reverts to Fig. 4A1; the faster neurons have a clear advantage whereas the slower neurons are much more likely to be suppressed which explains the suppression evident in the raster plot in Fig. 3A and the two rightmost heatmaps in Fig. 3C. The dynamics of the Type 1 population can be seen in the supplementary video sp1-20190619163238.mp4.

The phase portrait for minimal inhibition for type 2 in Fig. 4B1 shows that most neurons are again in a near threshold regime (as in Fig. 4 A1). Moreover, the average nullcline portrait is again close to a bifurcation, because the average fixed point requires only a small amount of additional excitation to move to the unstable branch of the V-nullcline as shown in Fig 1B. However, the vector flow near a subcritical Hopf bifurcation is completely different compared to the flow near a SNIC. For example, there is no slow channel. Instead, as explained in Fig. 2A, near a subcritical Hopf bifurcation, a bistable region exists in which quiescence at a stable fixed point (arrow) co-exists with repetitive spiking at the same value of net applied current. Whether the neuron is silent or active depends on recent history, meaning the current location of the trajectory. Specifically, the red closed curve in Fig. 4B1 is an unstable limit cycle that divides the neurons into two groups. Those inside the red curve will spiral into the stable fixed point at its center, whereas those outside will curve around it to fire an action potential. Thus the mechanism for cycle skipping has a component that results from the intrinsic dynamics. There is no bias toward faster cells, all neurons are almost equally likely to fall outside of the quiescent zone and fire an action potential. Fig. 4B2 shows that as neurons move to the right and fire actions potentials, inhibition accumulates as in Fig. 4A2 and again moves the V-nullcline downward, shifting the fixed point down and to the left. The unstable limit cycle opens up into a quasithreshold (not shown) that then merges with the unstable branch of the V-nullcline, labeled firing threshold. As in Fig. 4A3, Fig. 4B3 clearly shows two groups of neurons at the point of maximal inhibition, after all spiking neurons have fired on a given cycle. However, the group labeled skipped is still approaching their fixed points, and the distribution of these fixed points will be slanted rather than flat due to the sharper angle of the nnullcline at the intersection. The sharper slope is generic for type 2 versus type 1 because the steepness allow the n-nullcline to avoid the other branches of the V-nullcline, whereas multiple intersections are required for the SNIC in type 1. The dynamics of the type 2 population are shown in supplementary video sp2-20190619165631.mp4.

Fig. 4 C1 and C2 summarizes the movement of the stable fixed point due to changes in the level of inhibition. For type 1 in Fig. C1, the requirement that the n-nullcline be tangent to the V-nullcline at the SNIC bifurcation imposes a relatively flat slope on the n-nullcline near the intersection, which causes hyperpolarizing inhibition to move the fixed points horizontally, but not vertically (green arrows). The horizontal direction is the fast direction of the dynamics, therefore the trajec-tories remain on or near the n-nullcline with constant order. The firing threshold splits the population into spiking neurons on the right and suppressed skipping neurons on the left. Since the fixed points of the neurons are ordered with slower on the left and faster on the right, the movement of the fixed point does not alter their distribution. Therefore, slower neurons have a strong tendency to be suppressed with hyperpolarizing inhibition. In contrast, Fig. 4C shows the steeper slope of the n-nullcline required to switch the stability of the fixed point by moving between branches of the V-nullcline causes hyperpolarizing inhibition to move all the fixed points downward as well as leftward (green arrow). The trajectories in Fig. 4B3 for the skipping neurons do not follow the fixed point as Fig. 4A3, because the downward movement is in the slow direction. This downward and leftward trend moves most trajectories out of the unstable limit cycle when it forms in Fig. 4B1 around the rightmost fixed point in Fig. 4C2. This allows all neurons access to the fast curved vector fields that push them to an action potential trajectory, the composition of the neurons that spike on any given cycle is more evenly distributed throughout the population. This phenomenon has a contribution form anodal break excitation, also called post-inhibitory rebound, and explains why suppression is less prominent in the raster plot in Fig. 3B compared to Fig 3A, and the rightmost heatmaps of Fig. 3D compared to 3C.

### Steady State Synchrony in Oscillatory Networks- Shunting Inhibition

The results in the previous section were for hyperpolarizing inhibition. For both type 1 and type 2, the bifurcation that gives rise to spiking occurs at about −64 mV. Hyperpolarizing inhibition with a reversal potential of −75 mV hyperpolarizes the membrane at most points during the interspike interval except during the trough of the AHP. Shunting inhibition with a reversal potential of −65 mV does not produce big changes in the membrane potential. Fig. 5 shows the same simulations as Fig. 3 except for shunting versus hyperpolarizing inhibition. Cycle skipping to preserve population synchrony is still prominent in both types, as evidenced by the single neuron traces in the top left of parts A and B and by the peaks at integer multiples of the network frequency in the ISI histograms at lower right. The leftmost heatmaps(Fig. 5C1 and D1) confirm population synchrony is robust for both types, with generally lower participation (Fig. 5C2 and D2) than for hyperpolarizing inhibition at the same parameter values. In general, neurons that are not completely suppressed fire on average every other cycle. However, Type 2 networks clearly lose their superior resistance to suppression, as evidenced by the rasters in Fig. 5A versus 3B. Moreover, the histogram of participation for individual neurons is much flatter in the top right of Fig. 5A for type 2 with shunting inhibition compared to top right of Fig 3A for type two with hyperpolarizing inhibition, and in fact is slightly flatter than the histogram for type 1 with shunting inhibition in the top right of Fig. 5A. The two rightmost heatmaps once again show that these results are general, with CVs of participation that are slightly larger for type 2 (Fig. 5D3 vs C3), as well as more suppressed neurons at lower noise values for type 2 (Fig. 5D4 vs C4).

**Figure 5.**
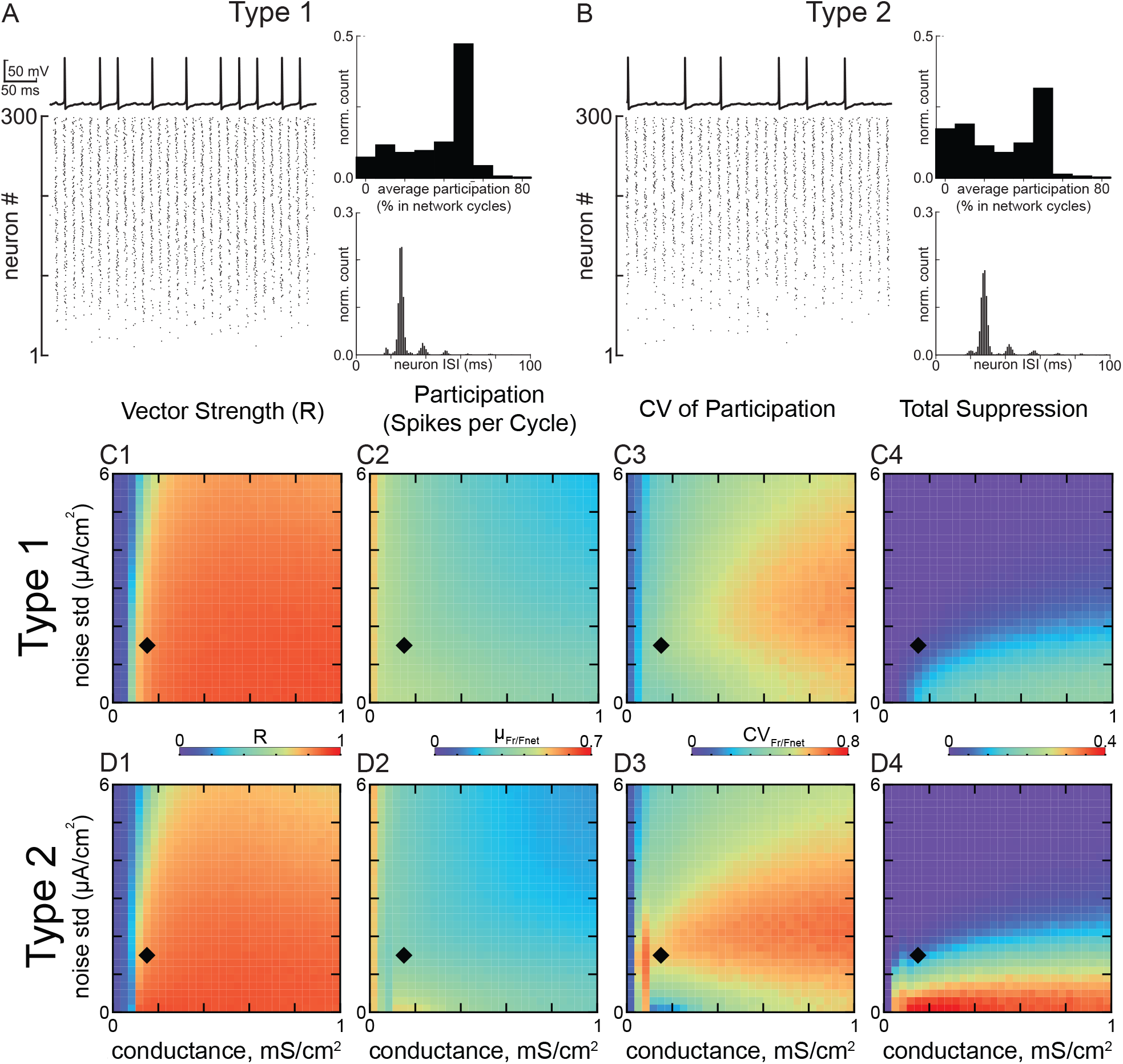
Steady State Synchronization of Heterogeneous Inhibitory Interneuronal Networks with Shunting Inhibition. A. Networks of type 1 neurons with synaptic conductance 1.5 mS/cm^2^ and noise standard deviation 1.5 µA/cm^2^, indicated by black diamonds in C and D. Bottom left: raster plot for 300 neurons, ordered by the level of applied depolarizing bias current, with the most depolarized neurons at the top. Top left: Representative membrane potential trace for an individual neuron that clearly exhibits a subthreshold oscillation due to network activity and cycle skipping. Bottom right: Histogram of interspike intervals across the population. Top right: Histogram of average participation for all neurons in the network. B. Networks of type 2 neurons with the same parameters as in A. The four panels are the same as in A. C. 2D parameter sweep in conductance strength and noise standard deviation for type 1 networks. Heatmaps from left to right give C1) the vector strength R of synchronization of individual spikes with the network oscillation, C2) the average participation for neurons that are not totally suppressed calculated as the mean µ_Fr/Fnet_ of the frequency of spiking neurons normalized by the population frequency, C3) the coefficient of variation of the participation CV_Fr/Fnet_, and C4) the fraction of completely suppressed neurons. D. 2D parameter sweep for type 2 networks. Heatmaps are the same as in C.

Figure 6 shows a phase plane analysis of the dynamics in a manner exactly analogous to Figure 4. There are two main differences. For type 1, in Fig. 6A the V-nullcline does not move as much with an inhibitory synaptic reversal potential of −65 mV as compared to −75 mV. Therefore, the fixed point moves very little (Fig. 6C1), keeping neurons that skipped near the firing threshold. The distribution of the dots representing neural trajectory are more compressed along the n-nullcline in Fig. 6A3, which reduces the advantage of the fastest neurons with the rightmost fixed points in escaping for the channel, hence the decrease in suppression. The V-nullclines and the corresponding fixed also move less with inhibition in Fig. 6B for type 2. One difference from Fig. 4B2 is that the quasithreshold described above (red curve in Fig. 6B2) is now visible as the “ghost” of the unstable limit cycle at the inhibition midpoint. Moreover, the inset in Fig. 6C2 reveals that the synaptic reversal potential of −65 mV is very close to the stable fixed point inside the red curve for the unstable limit cycle. A fraction of the population gets trapped inside the unstable limit cycle and skip a cycle. The slowest neurons tend to remain trapped. This explains the greater tendency for suppression in type 2 neurons for shunting versus hyperpolarizing inhibition. The movies corresponding to the phase portraits of type 1 and type 2 are found in sp3-20190619103132.mp4 and sp4-20190619104959.mp4, respectively.

**Figure 6.**
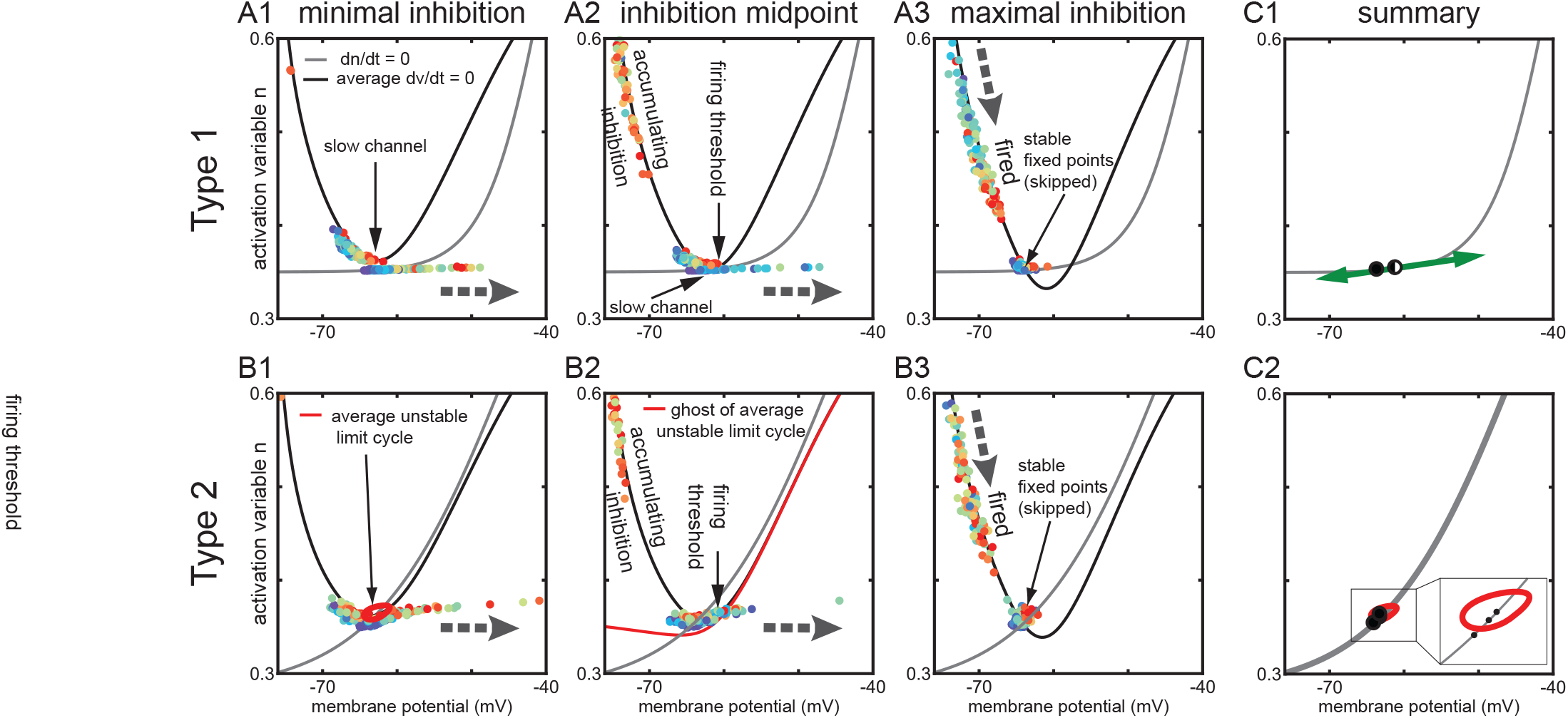
Phase Plane Analysis of Heterogeneous Inhibitory Interneuronal Networks with Shunting Inhibition. These phase plane portraits are similar to those in Fig. 4 except the inhibition is now shunting instead of hyperpolarizing. Large gray dashed arrows again show the local direction of motion in the vector field. Other thin black arrows simply point out a specific feature. The average V-nullcline (black curves) and the slow n-nullcine (gray) are shown for each type. Each again dot represents the current position of one neuron in the network, with the fastest neurons in red and the slowest in blue. A. Type 1. A1. Minimal inhibition again creates a slow channel between the nullclines (diagonal arrow). A2. Inhibition midpoint. The right branch of the V-nullcline is the firing threshold (vertical arrow). As neurons escape to the right (horizontal arrow shows direction of motion) and fire an action potential, inhibition again accumulates and moves the average V-nullcline down. A3. Maximal inhibition. The population is again segregated into two groups. However, the group that skipped this cycle (diagonal arrow) does not line up as clearly as in Fig. 4A along the flat portion of the n-nullcline. B. Type 2. B1. Minimal inhibition. In contrast to the absence of a fixed point in A1, there is a stable fixed point at the intersection of the average V-nullcline and the n-nullcine. This fixed point is again surrounded by an unstable limit cycle (red curve) that forms the boundary between spiking and quiescent trajectories. B2. Inhibition midpoint. As in A2 inhibition accumulates as neurons escape to the right and fire action potentials. However in this case the ghost of the unstable limit cycle unfurls into a quasithreshold (red curve) and separates spiking and skipping trajectories (vertical arrow labeling the red curve as the firing threshold). B3. Inhibition maximum. As in A3, two groups are evident corresponding to those that spiked and those that skipped (diagonal arrow). C. Summary. C1. This figure shows that there is less leftward movement of the fixed point as inhibition waxes and wanes in the fast, horizontal direction for type 1 compared to in Fig. 4C1. C2. The fixed point for type 2 tends to trap trajectories inside the unstable limit cycle (red curve, see blowup in inset), which enforces skipping.

### Robustness of the Mechanisms to Different Types of Heterogeneity and non-SNIC bifurcations

In Figs. 3 and 5, the applied current was varied to simulate heterogeneity in excitatory drive, as is typical [20, 36]. However, in addition to having different levels of excitatory input, different interneurons have different F/I curves. For type 1, the slope of the F/I curve is variable, and for type 2 neurons the cutoff frequency is also variable. We incorporated this additional dimension of variability into our simulations using a scale factor for the model dynamics as described in the Methods. In order to determine the effect of heterogeneity in this parameter, we scaled the dynamics of both variables for each neuron in order to achieve uniformly distributed cutoff frequencies in the range from 10 to 60Hz for type 2 neurons, and used the same range of scale factors for type 1 neurons. We then redid the heatmaps for each type with hyperpolarizing and with shunting inhibition (supplementary figures 1 and 2). Although both types show a reduction in synchronization, the qualitative results that type 2 networks are more resistant to partial and complete suppression for hyperpolarizing but not shunting inhibition still hold.

We also checked to make sure that a saddle-node-not-on-an-invariant-circle (SN) would have qualitatively similar synchronization tendencies. We adjusted the parameters 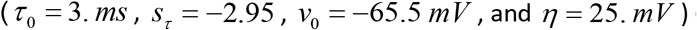 of the single type 1 model neuron such that the stable spiking limit cycle is not tangent to the saddle node at the bifurcation. Nonetheless, at synaptic conductance strengths sufficient to create a stable fixed point during network activity, a “bottleneck” or slow channel emerges with the same functional role in organizing network activity as if there were a saddle node on an invariant circle (not shown).

### Phase Amplitude Coupling

In the hippocampus, gamma power is maximal when nested in theta oscillations [37]. In order to determine the relative abilities of networks of neurons with type 1 excitability versus type 2 excitability to produce theta-nested gamma, we drove these networks with perfectly sinusoidal inhibitory waveforms at a fixed frequency in the theta range, in the presence of the constant heterogeneous depolarizing bias currents distributed as described in the Methods. The depolarization mimics tonic activation of metabotropic glutamatergic/cholinergic receptors. PV+ basket cells in freely moving rats fire at about 7 Hz during low oscillatory periods, but that rate triples to 21 Hz during theta oscillations [38], presumably due to greater tonic excitation. The sinusoidal drive mimics phasic inhibition from the septum. The amplitude of modulation is set to be the same for each neuron within the population, to reflect the common source of the modulation.

### Phase Amplitude Coupling- Hyperpolarizing Inhibition

Figure 7 gives examples of sinusoidal modulation at theta frequency of nested gamma oscillation. The time course of the modulation for both types of networks is the same and is given at the top of the figure. Panels A1,A2, B1 and B1 are have a representative single neuron trace at the top, a raster plot in the middle and a simulated LFP using the total inhibitory synaptic current summed across the network at the bottom. Both the raster lots and the simulated LFP show more neurons are recruited into the gamma rhythm in type 2 (Fig. 7B) networks compared to type 1 (Fig. 7A), both for a slower deeper modulation on the left, and a shallower faster modulation on the right. The noise and conductance parameters were selected in a regime in which all four types of networks synchronized well. The attributes of the network oscillations for the selected parameter regime are given in Table 3. In additional sets of simulations (not shown), we confirmed that the results shown below are qualitatively similar for wide range of synaptic conductance and noise levels and are not specific for the chosen values. Figure 7C shows the results in the two dimensional parameter space of modulation depth and modulation frequency, with the color in panel C1 and C2 indicating the un-normalized vector strength as described in the methods. If the vectors were normalized to reflect only how tightly locked the LFP envelope was to the theta drive, the two types perform equally. Removing the normalization reveals the greater recruitment of the population into the nested gamma in type 2 networks. The heatmaps in Fig. 7C1 and C2 show that low frequencies and shallow modulations are most effective, especially for type 2. The heatmap in Fig. 7D show that Type 2 always outperforms type 1 because the red color is always greater than zero.

**Figure 7.**
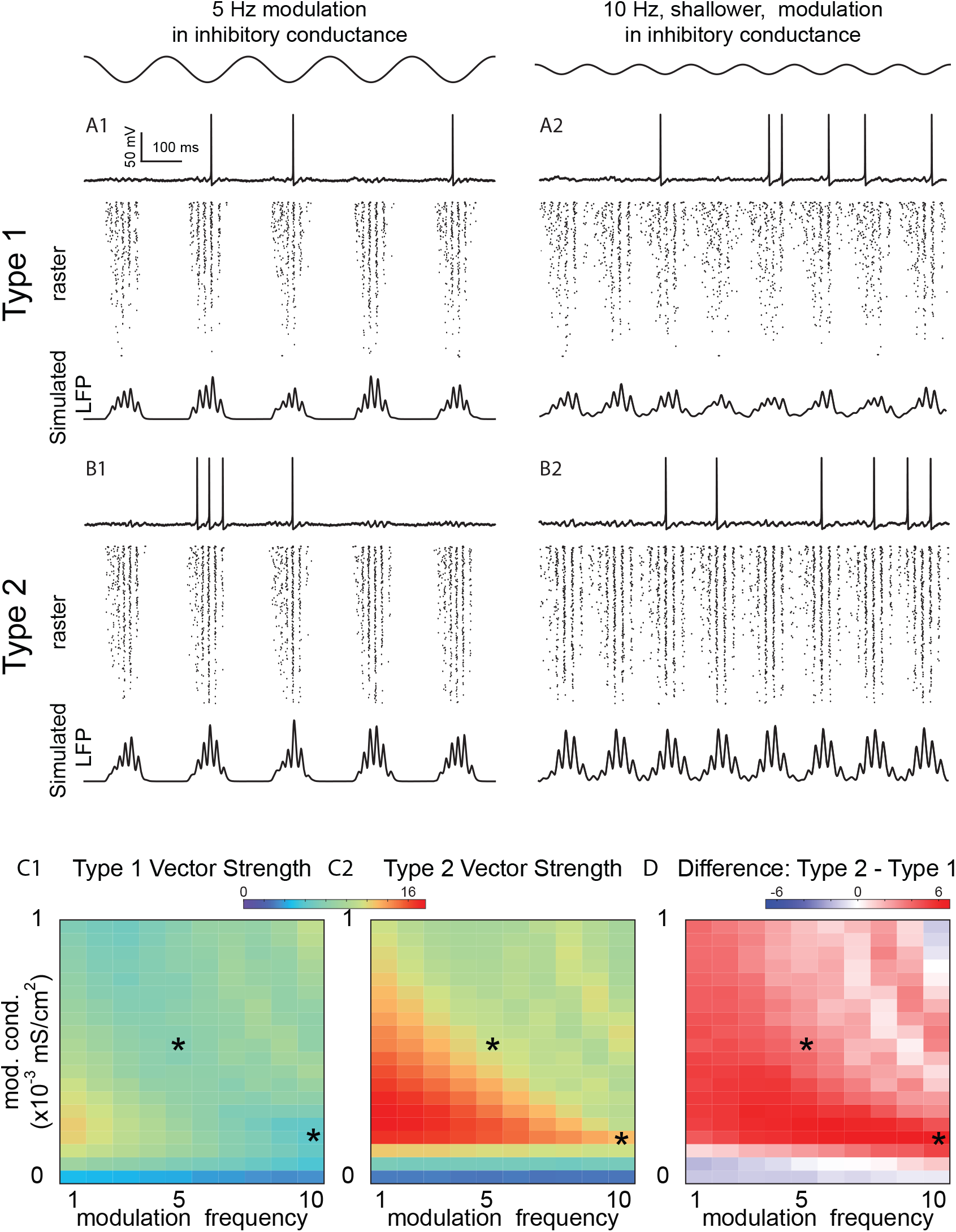
Theta Phase Gamma Amplitude Modulation of Heterogeneous Inhibitory Interneuronal Networks with Hyperpolarizing Inhibition. Two examples of sinusoidal modulation of these networks, with a 5 Hz sinusoidal modulation in inhibitory conductance shown at top left, and a shallower 10 Hz sinusoidal modulation shown at top right (conductance *g_ij_* = 0.1 *mS / cm*^2^, noise standard deviation *σ* = 3*μ A / cm*^2^). A. Type 1 networks. A1. 5 Hz modulation. A2. 10Hz modulation. Top: Representative sparsely firing single neuron type 1 traces with subthreshold oscillations due to network activity. Middle: Raster plots for 300 neurons with faster neurons (based on I app) shown at the top. Bottom: Simulated LFP consisting of summated inhibitory currents throughout the network. B. Type 2 networks. B1. 5 Hz modulation. B2. 10 Hz modulation. Top: Representative sparsely firing single neuron type 2 traces with subthreshold oscillations due to network activity. Middle: Raster plots for 300 neurons with faster neurons (based on I app) shown at the top. Bottom: Simulated LFP consisting of summated inhibitory currents throughout the network. C. Heatmaps for 2D parameter space of modulation depth (given in terms of the peak of the sinusoidal conductance waveform) and frequency. Asterisks show parameter values from A and B. C1. Vector strengths for type 1 networks. C2. Vector strengths for type 2 networks. C3. Difference of vector strengths for type 2 and type 1 networks. Hot colors indicate a difference greater than zero. The difference is well above zero in the lower lefthand cornerplot meaning that the vector strength for type 2 networks is higher.

**Table 3.** Characteristics of network steady-state oscillations with the parameters used in simulations with theta modulation: conductance g_ij_ = 0.1 mS / cm^2^, noise standard deviation *σ* = 3*μ* A / cm^2^.

|  | Hyperpolarizing Inhibition |  | Shunting Inhibition |  |
| --- | --- | --- | --- | --- |
|  | Type 1 | Type 2 | Type 1 | Type 2 |
| R | 0.8 | 0.88 | 0.75 | 0.67 |
| Mean participation | 0.2 | 0.27 | 0.22 | 0.17 |
| CV of participation | 0.81 | 0.64 | 0.64 | 0.65 |
| Total suppression | 0.15 | 0.03 | 0.04 | 0.04 |

Almost no theta modulation is evident in the membrane potential traces of individual neurons Fig 7A and B. As the network recovers from inhibitory modulation, the firing rate in the population increases, therefore network inhibition fills in when the external inhibitory drive wanes. This feedback mechanism is also responsible for keeping neurons near the bifurcation at minimal inhibition in steady-state oscillations (see Figs. 4A1 and B1 and 6A1 and B1).

In order to explain the superior performance of type 2 for hyperpolarizing inhibition, we can refer back to the phase plane in Fig. 4A. In order to synchronize, type 1 relies on the accumulation of inhibition as neurons escape through the slow channel, whereas there is no slow channel for type 2 (Fig. 4B). Instead, all neurons have access back to the strong vector fields away from the fixed point and an opportunity to spike. We hypothesize that the improved recruitment of neurons into the gamma oscillation in type 2 networks was because type 1 networks need more time to establish synchrony. We tested this idea in a network in which all neurons had the same applied bias current and therefore the same intrinsic frequency (see Figure 8A). The speed of synchronization is clearer for the homogeneous frequency case because participation is more uniform. A headto-head comparison over two cycles of sinusoidal revel reveals that indeed synchronization begins earlier in the type 2 homogeneous network compared to the type 1 homogeneous network (see leftmost vertical gray bar), which accounts in part for the larger vector strength for theta phase gamma amplitude coupling for type 2 with hyperpolarizing inhibition. The second gray bar shows that the synchrony persists longer for type 2 as well.

**Figure 8.**
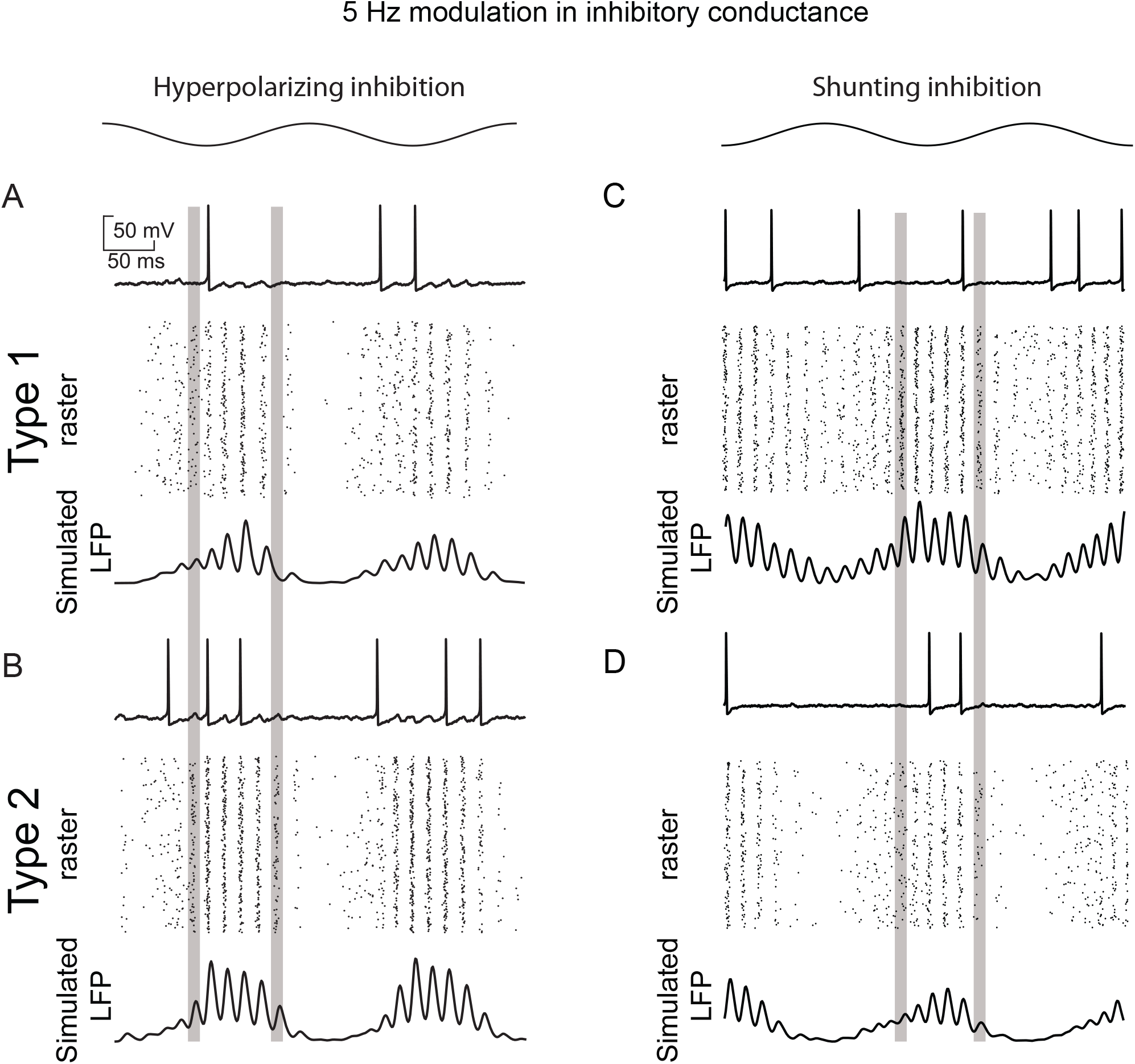
Speed of Recruitment of Theta-Nest Gamma in Homogeneous Inhibitory Networks with Hyperpolarizing Versus Shunting Inhibition. A. Homogeneous inhibitory network with hyperpolarizing inhibition showing that type 2 neurons are recruited more quickly into theta-nested gamma (leftmost vertical gray bar) and persist longer (rightmost gray bar). A1. Type 1. A2. Type 2. B. Homogeneous inhibitory network with hyperpolarizing inhibition showing that type 1 neurons are recruited more quickly into theta-nested gamma (leftmost vertical gray bar) and persist longer (rightmost gray bar). B1. Type 1. B2. Type 2.

The superiority of type 2 for hyperpolarizing inhibition derives from the ability to more quickly recruit the activity of sufficient neurons to establish synchrony at gamma frequencies. The presumption is that the interneurons are in an excited, oscillatory state during theta activity, and are rhythmically inhibited by the septum. For modulation that is too shallow, gamma activity is ongoing and theta power is weak (bottom blue strips on Fig.7 C1 and C2). For modulation that is sufficiently deep to establish theta power, the modulation must be sufficiently slow or shallow that there are still long enough windows of time above the spiking threshold to recruit enough active neurons to establish gamma synchrony (red areas in bottom left half of Fig. 7 C1 and C2).

### Phase Amplitude Coupling-Shunting Inhibition

The faster recruitment of type 2 oscillators did not persist for homogeneous networks with shunting inhibition (Fig. 8B). Instead, type 1 oscillators are recruited more quickly into nested theta gamma because shunting inhibition clamps neurons closer to the spiking threshold (see Fig. 6C1 vs. Fig. 4C1). Type 2 networks clearly lose their superiority, likely because of the tendency of type 2 oscillators with shunting inhibition to remain trapped at the fixed point inside the unstable limit cycle (Figure 6B1 and C2).

Figure 9A and B show examples of Type 1 and 2 networks with shunting inhibition. Just as switching from hyperpolarizing to shunting inhibition nullified the advantage of type network in robustness to suppression during steady state synchrony shown in Figures 3 and 5, changing the inhibition from hyperpolarizing to shunting nullifies the advantage of type 2 for phase amplitude coupling as predicted by Fig. 8. In fact, Fig. 9D shows that type 1 networks have better phase amplitude coupling for deep modulations (blue indicates a type 2 type 1 difference is less than zero). Although type 2 appears to perform better at low modulation amplitudes, the vector strength is so small (blue at the bottom of Fig. 9C1 and C2) as to render this regime unimportant. The tendency of the unstable limit cycle in Fig. 6B1 to trap trajectories clearly reduces the speed of synchronization of type 2, and is likely responsible for its poorer performance with shunting inhibition. Supplementary figures 3 and 4 shows that the PAC tendencies for type 1 versus type 2 are preserved under the second type of heterogeneity utilized in supplementary figures 1 and 2.

**Figure 9.**
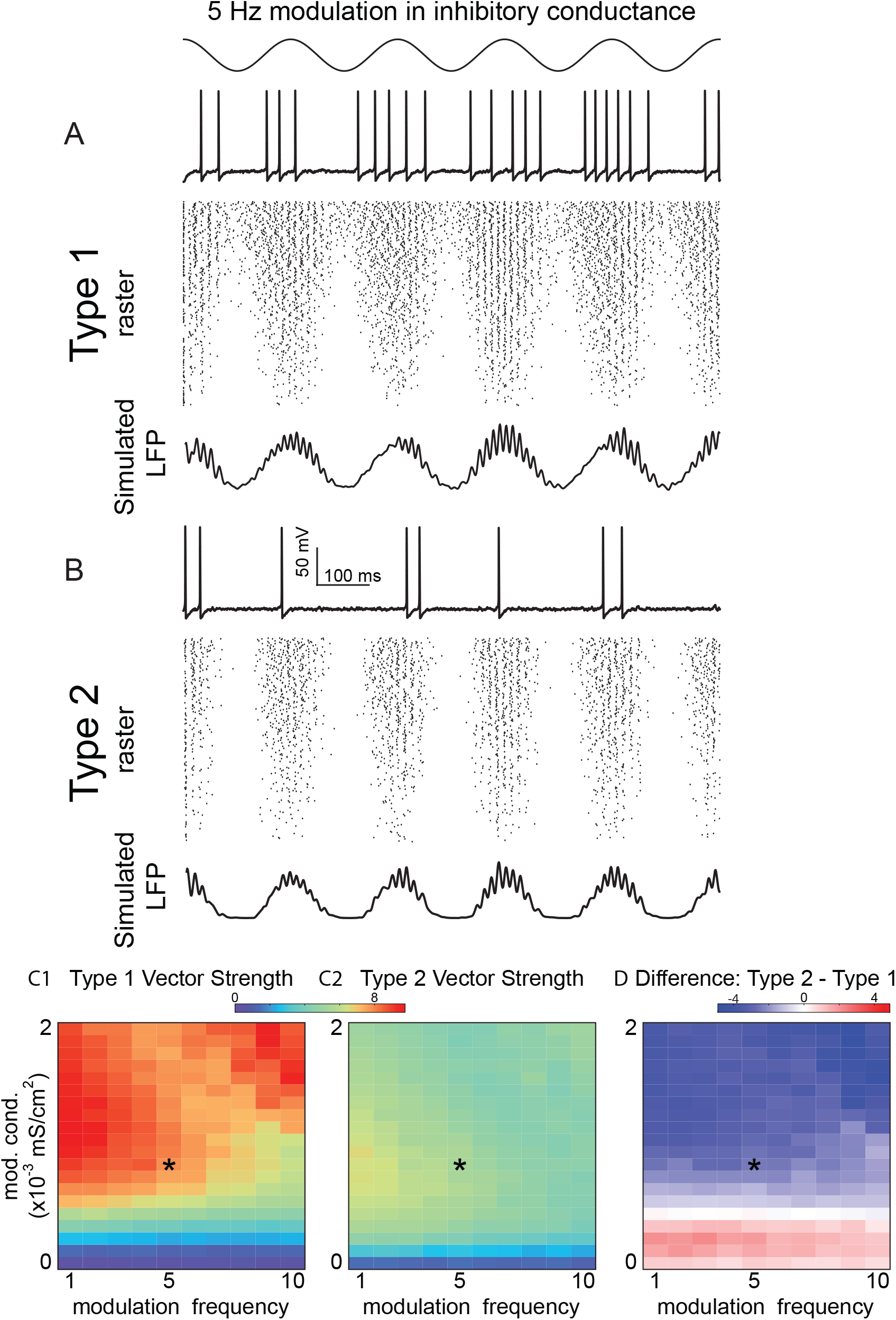
Theta Phase Gamma Amplitude Modulation of Heterogeneous Inhibitory Interneuronal Networks with Shunting Inhibition. One example of sinusoidal modulation of these networks, with a 5 Hz sinusoidal modulation in inhibitory conductance shown at the top for the same synaptic conductance and noise standard deviation as in Fig.7. A. Type 1 networks modulation. Top: Representative sparsely firing single neuron type 1 traces with subthreshold oscillations due to network activity. Middle: Raster plots for 300 neurons with faster neurons (based on I app) shown at the top. Bottom: Simulated LFP consisting of summated inhibitory currents throughout the network. B. Type 2 networks. Top: Representative sparsely firing single neuron type 2 traces with subthreshold oscillations due to network activity. Middle:. Raster plots for 300 neurons with faster neurons (based on I app) shown at the top. Bottom: Simulated LFP consisting of summated inhibitory currents throughout the network. C. Heatmaps for 2D parameter space of modulation depth (given in terms of the peak of the sinusoidal conductance waveform)and frequency. Asterisk shows the parameter values from A and B. C1. Vector strengths for type 1 networks. C2. Vector strengths for type 2 networks. C3. Difference of vector strengths for type 2 versus type 1 networks. Cool colors indicate a difference less than zero. The difference is below zero in this plot, meaning that the vector strength for type 2 networks is higher, except at the bottom where the vector strength is relatively small.

## DISCUSSION

### Summary of Results

We present two major results. The first is that networks of heterogeneous neural oscillators with type 2 excitability are more resistant to suppression of the slower oscillators than those with type 1 excitability when they are coupled with hyperpolarizing inhibition but not with shunting inhibition. The second result is that theta phase to gamma amplitude coupling is more strongly recruited in the type 2 networks, again for hyperpolarizing but not shunting inhibition. Moreover, we provide mechanistic explanations for these phenomena, which frame conditions for the generality of these results.

The tendency of slower type 1 neurons to be more suppressed than type 2 neurons for hyperpolarizing inhibition relies on two principles. The first is the creation by inhibition of a slow channel near the SNIC bifurcation for type 1. The channel serves to line the fixed points of the heterogeneous oscillators up along the slow nullcline and along the direction of motion of the fast subsystem. This arrangement clearly favors the faster oscillators, and is less prominent for shunting versus hyperpolarizing inhibition.

The second principle is the circular motion about the subcritical Hopf bifurcation for type 2 neurons and the existence of a boundary in the phase plane between spiking and skipping neurons allows for broader participation in steady state network oscillation for hyperpolarizing but not shunting inhibition. The critical assumption is that the stable fixed point just prior to the Hopf bifurcation lies near the shunt reversal potential, but well above the hyperpolarizing reversal potential. This assumption seems reasonable based on the definitions of hyperpolarizing and shunting inhibition combined with the location of the Hopf at the point that positive feedback from the sodium channel destabilizes the rest potential. Thus hyperpolarizing inhibition pushes all neurons leftward (in the fast direction) and the circular motion brings them downward and into the fast vector fields that give neurons a much more equal chance to fire than type 1. However, shunting inhibition, rather than freeing trajectories from the unstable limit cycle surround the stable fixed point just prior to the Hopf, traps them within that limit cycle, favoring suppression.

These principles apply to the transient synchronization in theta-nested gamma as follows. Note that the steady state synchronization and participation are similar between type 1 and type 2. However, type 2 neurons are more quickly recruited into nested theta gamma for hyperpolarizing inhibition because their intrinsic dynamics do not require accumulation of synaptic inhibition to split the populations into spiking and skipping groups. Faster recruitment also allows synchronization, which requires a minimum number of active neurons, to persist longer. The phase plane portrait also accounts for the slower recruitment of type 2 neurons with shunting inhibition into nested theta gamma and the shorter persistence that accompanies late recruitment. The mechanism is again the tendency of shunting inhibition to clamp the type 2 trajectories inside the unstable limit cycle, which slows recruitment into theta nested gamma.

### Generality

A heterogeneous, multicompartmental neuron can be considered as a chain of diffusively coupled oscillators. Although the SNIC bifurcation is generic in single compartment models, studies of heterogeneous oscillators, each exhibiting a SNIC in isolation [39], exhibit a complicated solution structure in a diffusively coupled system. It is not clear that a heterogeneous, multicompartmental neuron can exhibit a SNIC precisely, so it was important to show that the principles described in the preceding section for type 1 neurons still hold for a network in which individual neurons do not have a SNIC, but rather a SN not on an invariant limit cycle. The slow channel still forms.

Another important way in which we checked the generality of the results was to introduce a second type of heterogeneity. We varied the slope of the f/I curve for type 1 neurons and both the cutoff frequency and slope for the type2 neurons by rescaling the temporal dynamics. The cutoff frequency for fast spiking PV+ basket cells [40] had an estimated standard deviation of 40%. Thus, the 10-60 Hz range in cutoff frequencies explored should map onto biologically plausible distributions of cutoff frequencies. Qualitatively, our major results were preserved under these manipulations (Supplementary Figs. 1-4).

### Previous comparisons of type 1 versus type 2 excitability

Previously Rinzel and Ermentrout [41] contrasted the phase portraits and bifurcation structure for type 1 versus type 2 excitability using the Morris Lecar [42] model in the two regimes. Izhikevich [16] also showed in the phase plane that various minimal conductance-based models could exhibit saddle node and Hopf bifurcations. Neither study explored the implications of the bifurcation type for synchronization.

In the weak coupling regime, Ermentrout and colleages [43, 44] found that type 2 neurons receiving noisy common input synchronize better than type 1. Borgers and Walker [45] compared the synchronization tendencies of type 1 [20] and type 2 [46] interneurons embedded in excitatory/inhibitory networks with type 1 excitatory neurons. They found that when gap junctions were incorporated into their model, there was a sharper transition from pyramidal interneuronal network gamma to interneuronal network gamma in the type 2 networks compared to type 1 networks. Rich et al [47] found that excitatory/inhibitory networks with type 2 interneurons were more robust to changes in network connectivity compared to type 1. None of the type 1 versus type 2 model comparisons cited above were calibrated to match spike shape, shape of the F/I curve, time constant and input resistance across excitability types as were the models in our study. Thus, we extend these previous comparisons so that any difference in network activity is due to the bifurcation type.

### Cycle skipping mechanisms

Cycle skipping to enforce population synchrony within the inhibitory neuron population has been observed in several studies in networks with both type 1 [48], and type 2 interneurons [17, 49]. However, the bifurcation structure for the four cases (combinations of type 1 and type 2 excitability with either hyperpolarizing or shunting inhibition) has not been examined in detail previously. It is clear that interneurons do not fire on every gamma cycle *in vivo*. For example, CA3 FS cells fired at 21 Hz during 45 Hz gamma at 45 Hz, thus they fired roughly on every other cycle [50]. Cycle skipping provides a mechanism by which coupled oscillator models can produce tightly synchronized firing with sparse firing in individual neurons. This mechanism is distinct from a clustering mechanism that was advanced to introduce robustness of population synchrony. In that mechanism, interneurons did not participate on every gamma cycle because synchronized subclusters fired in sequence [51]. Previously [17], we found that cycle skipping more prominent in Type 2 networks, but it was not possible to do a fair comparision across the parameter space with the unmatched models we used in that study. The appropriate conclusion is that cycle skipping is intrinsic to the dynamics of type 2 neurons connected by hyperpolarizing inhibition, but in all other cases synaptic mechanisms predominate over intrinsic mechanisms. Cycle skipping allows robustness to heterogeneity in excitatory drive and connectivity for both type 1 and type 2 networks, overcoming a perceived weakness of coupled oscillator models of network gamma since they were introduced (see next section).

### Previous studies on suppression

Computational studies have often shown that fast oscillations based on reciprocal inhibition are exquisitely sensitive to heterogeneities in the network [1,20,23,36]. Inhibitory neural networks generally lose synchrony as heterogeneity is increased in one of two ways, phase dispersion or suppression, depending on the ratio of the time constant for decay of inhibition to the population frequency [36, 52]. Here, we looked at fast synapses, which tend to favor the suppression regime over phase dispersion. Early work [20,36,52] on robustness of inhibitory interneuronal synchrony focused of type 1 excitability only. These studies did not include conduction delays, and concluded that hyperpolarizing inhibition and non-physiological low levels of heterogeneity in excitatory drive were required for synchrony. Subsequent studies [25] used the same type 1 Wang and Buzsaki model added a ring topography with conduction delays ranged from 0.7 to 10.5 ms; they found that networks with shunting inhibition were more robust to heterogeneity than with hyperpolarizing inhibition. Their raster plots clearly show more suppression for type 1 for hyperpolarizing compared to shunting inhibition, in agreement with our findings. They repeated their simulation with the type 2 model of Erisir et. al. [46], and obtained similar synchronization results, as in our study. However, they did not show raster plots or report on suppression in the type 2 networks. We also used a distribution of delays (from 0.7 to 3.5) to prevent the formation of two cluster solutions [35] and stabilize global synchrony by moving the operating point away from the destabilizing discontinuity in the phase resetting curve at 0 and 1.

### Therapeutic directions for PAC and cognition

The hippocampal theta rhythm has been shown to be necessary for spatial learning by rats in a water maze [53]. Phase amplitude coupling between theta and gamma likely plays an important role in cognition [54], and we have shown here that type 2 excitability in interneurons strengthens phase amplitude coupling between theta and gamma rhythms for hyperpolarizing but weakens it for shunting inhibition. This suggests the possibility that interneurons could modulate their excitability type according to whether inhibition is hyperpolarizing or shunting, or vice versa. Moreover, this suggests a therapeutic strategy to manipulate excitability type, and thereby theta nested gamma phase amplitude coupling, by targeting currents active in fast spiking interneurons in the subthreshold regime to tip the balance toward outward currents.

The balance of inward and outward currents in the voltage range traversed between action potentials during the interspike interval (ISI) determines whether a neuron exhibits type 1 or type 2 excitability, and is easily modifiable by altering this balance [55, 56]. For type 2, outward currents predominate at steady state in this region of subthreshold membrane potential, but are activated more slowly than the inward currents. Therefore, the dwell time in this region of membrane must be brief so that the outward current does not equilibrate and stop the depolarization caused by the inward currents between spikes. There is a maximum ISI for which repetitive spiking can be supported, hence a minimum frequency below which the neuron cannot sustain repetitive firing. In contrast, inward currents predominate at steady state in the subthreshold range on membrane potentials spanned by the interspike interval, therefore arbitrarily low firing rates can be sustained. Decreasing inward or increasing outward currents that have a steady state component during the ISI favors type 2 excitability, and the opposite manipulations favor type 1. These manipulations, depending upon whether inhibition is hyperpolarizing or shunting, could increase theta/gamma PAC. Increased theta/gamma phase amplitude coupling may in turn improve some aspects of cognition.

## Detailed Methods

### Network

For all simulations we used 300 neurons of the same excitability type, connected by bi-exponential inhibitory synapses. In the network, the input current for each neuron is given by the following equations:

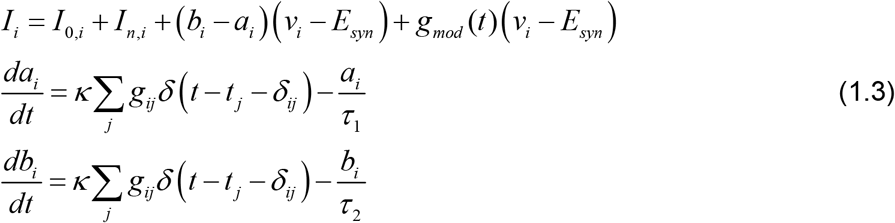

where *v_i_* and *I_i_* are membrane potential and input current of i^th^ neuron; *I_0,i_* is an applied current; 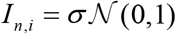 is a noise current with an independent random process with zero mean and unit variance for each neuron 𝒩 (0,1). These processes were sampled every 0.1 ms, and the current was linearly interpolated between these times to produce consistent results regardless of the time step. The synaptic rise and fall time constants were 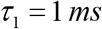 and 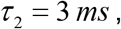 respectively. The synaptic reversal potential *E_syn_* was set to −75 mV for hyperolarizing and −65mV for shunting inhibition. The conductance (*g_ij_*) and conduction delay (*δ_ij_*) between the *i^th^* and *j^th^* neurons are given in units of mS/cm^2^ and ms respectively; *δ* () is Dirac’s delta function; and ***k*** is a normalization constant to keep peak of *b_i_*-*a_i_* for unitary PSG equal to *g_ij_*

For both steady-state and sinusoidally modulated network oscillations, *I_0,i_* were drawn from uniform distribution with the range [2, 3.8] *μA / cm*^2^ providing a distribution of intrinsic frequencies with a 20 Hz range (see vertical and horizontal bars Figure 2A). In steady-state oscillations regime inhibitory modulatory conductance *g_mod_* was set to zero. In a contrast, for modulated network oscillations *g_mod_* was sinusoidally modulated: 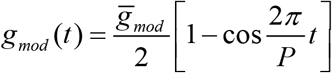, where P is the period, and 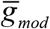 is the amplitude of modulation.

Connections in the network were sparse and random with probability *p* = 0.133 of connection between any given the *i^th^* and *j^th^* neurons. For all simulations presented here, conduction delays were uniformly randomly distributed between 0.7 and 3.5 ms.

### Numerical simulations and bifurcation analysis

The bifurcation analysis was performed in XPPAUTO [57].

The network models were implemented as Python 2.7 script for the simulation package NEURON [58]. The integration time step was constant at 0.01 ms. Synaptic activation initially was set to zero for all simulations. The membrane potential of each neuron was initialized randomly from a normally distribution with mean −50mV and standard deviation of 20 mV and the slow n variable was initialized at the steady-state value for the membrane potential. The data presented on vector strength, participation, CV of participation and total suppression for networks biased in the oscillatory regime were averaged over ten trials at each parameter setting for runs of duration 2.5 s with the initial 500 ms ignored to minimize the effects of transients. Each trial had its own random connectivity pattern, random initialization of the state variables, random distribution of bias currents, random delay distribution and random noise sources. For the sinusoidal drive simulations the PAC was averaged over 20 periods of sinusoidal drive, again averaged over 10 trials as described above, but no transients were deleted. Simulations were run on Louisiana Optical Network Infrastructure, QB2-cluster. The source code is available at https://senselab.med.yale.edu/ModelDB/showmodel.cshtml?model=259366#tabs-1

(will be publicly accessible upon publication; code for reviewers ZhD7rUh64oD1roKUHc7D). The phase of each spike within a cycle used was calculated using the length of that particular cycle. Cycle lengths are variable and were computed using the peaks in the population rate as described in Tikidji-Hamburyan et al. [17]. The phase was used to construct the vectors for the vector strength calculation.

In the supplementary figures, we applied a multiplicative scale factor (*F_i_*) to the rates of change of both variables. Since the Gaussian noise term simulates Brownian motion in the membrane potential, in which distance is proportional to the square root of time, the noise term as then divided by the square root of the scale factor. Taken together, these manipulations simply scale the intrinsic frequency since all intrinsic (but not synaptic) processes are sped up or slowed down equally.

### Measure of phase amplitude coupling

Since we apply the theta drive, there is no uncertainty with respect to the phase of the theta oscillation, in contrast to the uncertainty in experimental data such as local field potentials or the EEG. This simplified our analysis. Moreover, since we apply the exact same amplitude of theta modulation to different networks, we were interested not only in the tightness of the coupling of the theta drive and the evoked nested gamma oscillation, but also in the magnitude of the gamma oscillations. Therefore in order to quantify the coupling between theta phase and gamma amplitude, we choose the mean vector length (MVL) [59, 60], but without normalization of the amplitude. The vectors consisted of the known theta phase with the magnitude given by the amplitude of the gamma envelope determined using the Hilbert transform of the simulated LFP. The simulated LFP was the synaptic inhibitory current summed over the network. The sum of these vectors produces a vector strength that is not bounded between 0 and 1, but which does accurately reflect the amplitude of the nested gamma oscillations evoked by a constant sinusoidal stimulus at theta frequency, and the preferred theta phase. The normalized vector strength shows only how strongly the gamma envelope is locked to the preferred theta phase. The unnormalized version takes into account the actual amplitude of the gamma envelope.

---

**Supplementary Figure 1. Steady State Synchronization of Heterogeneous Inhibitory Interneuronal Networks with Hyperpolarizing Inhibition with Additional Heterogeneity in Intrinsic Dynamics.** In addition to a uniform distribution of applied currents *I_0,i_ =* [*2, 3.7*] *μ A / cm^2^*, heterogeneity in *F_i_* = 1.04 ± 0.4 normally was introduced in the population. *F_i_* gives a range of cutoff frequencies from 10 to 65 Hz for type 2. 2D parameter sweep in conductance strength and noise standard deviation for type 1 networks. Heatmaps from left to right: A1: vector strength R of synchronization of individual spikes with the network oscillation, A2: average participation for neurons that are not totally suppressed calculated as the mean µ_Fr/Fnet_ of the frequency of spiking neurons normalized by the population frequency, A3: the coefficient of variation of the participation CV_Fr/Fnet_, and A4: the fraction of completely suppressed neurons. B. 2D parameter sweep for type 2 networks. Heatmaps are the same as in A. The qualitative result that there is more total suppression in type 1 networks is preserved. Type 2 networks are still able to achieve stronger R > 0.9 synchronization almost everywhere, whereas Type 1 networks can reach *R* ≈ 0.85 in small subregion and show moderate synchronization *R* ∈ [0.6, 0.8] in the rest of parameter space. In additional, heterogeneity in the neuron dynamics reduces average participation in network oscillation and dramatically increases variability in participation for Type 1 networks (*CV* from 0.8 to 1.0 almost everywhere) but moderate increase in area but not in maximum value (max *CV* ≈ 0.85) for Type 2networks. Finally, both types show increase in total suppression.

**Supplementary Figure 2. Steady State Synchronization of Heterogeneous Inhibitory Interneuronal Networks with Shunting Inhibition with Additional Heterogeneity in Intrinsic Dynamics. 2D parameter sweep in conductance strength and noise standard deviation for type 1 networks.** Heatmaps from left to right: A1: vector strength R of synchronization of individual spikes with the network oscillation, A2: average participation for neurons that are not totally suppressed calculated as the mean µ_Fr/Fnet_ of the frequency of spiking neurons normalized by the population frequency,A3: the coefficient of variation of the participation CV_Fr/Fnet_, and A4: the fraction of completely suppressed neurons. B. 2D parameter sweep for type 2 networks.

Heatmaps are the same as in A. The qualitative result that there is more total suppression in type 2 networks is preserved. The additional heterogeneity is the same as in Supplementary Figure 1.

**Supplementary Figure 3. Theta Phase Gamma Amplitude Modulation of Heterogeneous Inhibitory Interneuronal Networks with Hyperpolarizing Inhibition with Additional Heterogeneity in Intrinsic Dynamics.** Heatmaps for 2D parameter space of modulation depth and frequency. A. Vector strengths for type 1 networks. B. Vector strengths for type 2 networks. C. Ratio of vector strengths for type 2 versus type 1 networks. Cool colors indicate a ratio less than one. The ratio is always above one in this plot meaning that the vector strength for type 2 networks is higher, as in Figure 7. The additional heterogeneity is the same as in Supplementary Figure 1.

**Supplementary Figure 4. Theta Phase Gamma Amplitude Modulation of Heterogeneous Inhibitory Interneuronal Networks with Hyperpolarizing Inhibition with Additional Heterogeneity in Intrinsic Dynamics.** Heatmaps for 2D parameter space of modulation depth and frequency. A. Vector strengths for type 1 networks. B. Vector strengths for type 2 networks. C. Ratio of vector strengths for type 2 versus type 1 networks. Cool colors indicate a ratio less than one. The ratio is always below one in this plot as in Figure 9, meaning that the vector strength for type 2 networks is higher, except at the bottom where the vector strength is so small that there is negligible PAC. The additional heterogeneity is the same as in Supplementary Figure 1.

**Supplementary movie 1 (sp1-20190619163238.mp4) Dynamics of the Type 1 population with hyperpolarizing inhibition.** Left plots from top to bottom: population raster plot: Color code indicates amplitude of excitatory drive; Population voltage traces; population slow variable traces; averaged synaptic conductance of the population. Right: phase plane analysis. Position of each neuron in the state space is marked by a color cycle. Color code is the same as in top left plot. Blue solid line indicates average instantaneous voltage nullcline. Dashed blue line indicates voltage nullcline in the absence of inhibition. Red dashed line shows slow variable nullcline.

**Supplementary movie 2 (sp2-20190619165631.mp4) Dynamics of the Type 2 population with hyperpolarizing inhibition.** Plots and lines are the same as for Supplementary movie 1. An additional black dash-dot curve indicates unstable limit cycle which breaks and turns in quasithreshold.

**Supplementary movie 3 (sp3-20190619103132.mp4) Dynamics of the Type 1 population with shunting inhibition.** Plots and lines are the same as for Supplementary movie 1.

**Supplementary movie 3 (sp4-20190619104959.mp4) Dynamics of the Type 2 population with shunting inhibition.** Plots and lines are the same as for Supplementary movie 2.

